# Sentinel *p16*^*INK4a*^+ cells in the basement membrane form a reparative niche in the lung

**DOI:** 10.1101/2020.06.10.142893

**Authors:** Nabora Reyes de Mochel, Ka Neng Cheong, Monica Cassandras, Chaoqun Wang, Maria Krasilnikov, Peri Matatia, Ari Molofsky, Judith Campisi, Tien Peng

## Abstract

Senescent cells are recognized drivers of aging-related decline in organ function, but deciphering the biology of senescence *in vivo* has been hindered by the paucity of tools to track and isolate senescent cells in tissues^1–4^. Deleting senescent cells from transgenic murine models have demonstrated therapeutic benefits in numerous age-related diseases^5–11^, but the identity, behavior, and function of the senescent cells deleted *in vivo* remain elusive. We engineered an ultra-sensitive reporter of *p16*^*INK4a*^, a biomarker of senescence^12^, to isolate and track *p16*^*INK4a*^+ cells *in vivo*. Surprisingly, *p16*^*INK4a*^+ mesenchymal cells appear in the basement membrane adjacent to epithelial progenitors in the lung shortly after birth, and these cells demonstrate senescent characteristics *in vivo* and *ex vivo*. Transcriptomic analysis of *p16*^*INK4a*^+ mesenchymal cells from non-aged lungs demonstrates a transition to a secretory phenotype upon airway epithelial injury. Heterotypic 3D organoid assays show that injured *p16*^*INK4a*^+ mesenchymal cells enhance epithelial progenitor proliferation, and we identified EREG as a novel airway progenitor mitogen produced by the secretory *p16*^*INK4a*^+ mesenchymal cells. Mesenchymal-specific deletion of the *p16*^*INK4a*^ gene abrogates features of senescence *in vivo*, but also attenuates normal epithelial repair. Thus, *p16*^*INK4a*^+ mesenchymal cells can act as sentinels for the airway epithelial stem cell niche, poised to transition to a senescence-associated secretory phenotype to support barrier repair. Our data identify possible cellular targets *in vivo* for a rapidly growing list of senolytic therapies, but also raises important questions about the hidden cost of targeting senescent cells present in normal organs.

## Main

p16^INK4a^ is a tumor suppressor encoded in the *Cdkn2a* locus that is upregulated in cultured cells *in vitro* undergoing cellular senescence^13^, a form of cell cycle arrest associated with aging. Mouse reporters using *p16*^*INK4a*^ promoter to drive luciferase expression have consistently demonstrated upregulation of *p16*^*INK4a*^ with aging and wound repair^14,15^. Based on these and other studies quantifying *p16*^*INK4a*^ transcripts in whole tissues, the prevailing idea is that *p16*^*INK4a*^+ cells are rare or absent in young and healthy tissues, suggesting that senescence does not play a role in normal tissue maintenance. However, the use of luciferase combined with whole-body bioluminescence imaging precludes studying *p16*^*INK4a*^+ cells at the cellular resolution. Genetic models expressing suicide genes have demonstrated beneficial effects of killing *p16*^*INK4a*^+ cells in numerous models of aging-related pathologies, but the identity and behavior of living *p16*^*INK4a*^+ cells in their cellular ecosystem within tissues remain largely undefined. The lack of clear definition of living *p16*^*INK4a*^+ cells and their function hinder the study of senescence *in vivo*, and impedes the design of rational therapeutics targeting these cells.

### High-sensitivity fluorescent reporter of senescence *in vivo*

We first set out to define *p16*^*INK4a*^+ cells that might play a role in lung injury repair using the existing 3MR mouse expressing mRFP (monomeric red fluorescent protein) in addition to luciferase and a viral thymidine kinase driven by the *p16*^*INK4a*^ promoter^16^. Utilizing the naphthalene injury model where airway epithelial progenitors encounter replicative stress to repair the damaged airway epithelium^17,18^, we show that the *p16*^*INK4a*^ transcript is upregulated in the whole lung after injury (Extended Data Fig. 1a). However, we failed to detect RFP+ cells in the naphthalene-injured lungs of the 3MR animals (Extended Data Fig. 1b), despite our qPCR showing an increase in *p16*^*INK4a*^ transcript after injury. We postulated that the failure to detect fluorescent *p16*^*INK4a*^+ cells in this setting might be due to: 1) the relative low abundance of the *p16*^*INK4a*^ transcript in cells in which the *p16*^*INK4a*^ promoter is active, and 2) the low fluorescence intensity and/or stability of mRFP. Our solution was to construct a bacterial artificial chromosome (BAC) where tandem cassettes of H2B-GFP is expressed in frame with the *p16*^*INK4a*^ gene product in the murine *Cdkn2a* locus, thus utilizing *p16*^*INK4a*^ promoter to drive the expression of multiple copies of a stable fluorescent protein that would be incorporated into the nucleosome (Extended Data Fig. 1c, Method for targeting strategy). The BAC was injected into embryos to create a transgenic model named the <u>INK</u>4A H2<u>B</u>-GFP <u>R</u>eporter-<u>I</u>n-<u>T</u>and<u>e</u>m (hereafter referred to as *INKBRITE*) mouse. Flow cytometry analysis of naphthalene injured *INKBRITE* lungs showed an increase in highly fluorescent GFP+ cells compared to uninjured *INKBRITE* lungs (Fig. 1a, and Extended Data Fig. 1d), and isolated GFP+ cells shows significant upregulation of the *p16*^*INK4a*^ transcript (Fig. 1a).

**Figure 1.**
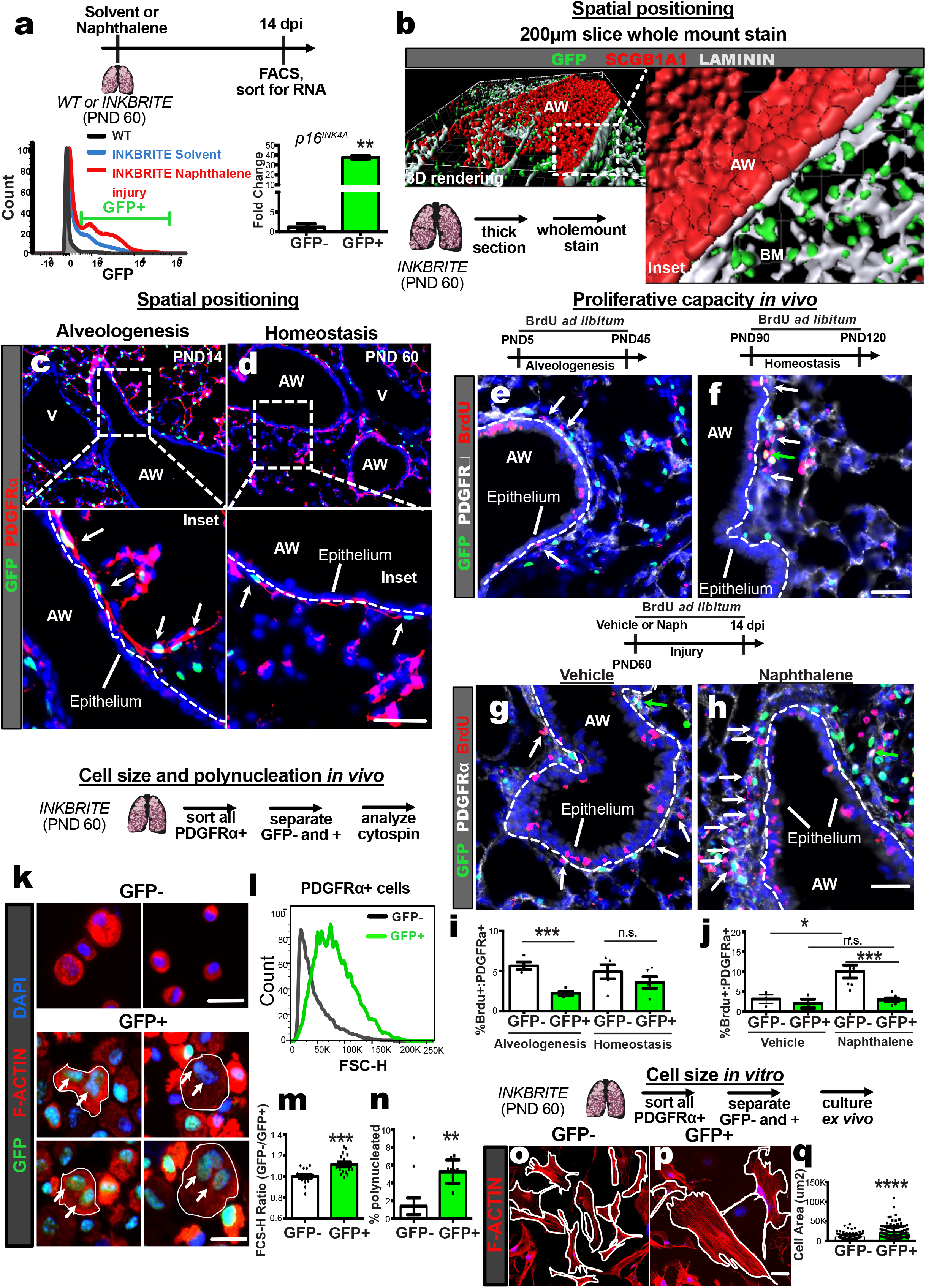
*INKBRITE* identifies *p16*^*INK4a*^+ cells with senescent characteristics *in vivo*. **a,** FACS analysis of GFP from *INKBRITE* lungs and qPCR of sorted GFP+ and – cells. (n=4) **b,** Wholemount image of thick-sectioned *INKBRITE* lung rendered on Imaris. **c,d,** Immunohistochemistry (IHC) of GFP overlaid with PDGFRα during alveologenesis (PND14) and homeostasis (PND60) phases of postnatal lung maturation. **e-h,** IHC of BrdU incorporated into PDGFRα+/GFP− (white arrows) or PDGFRα+/GFP+ (green arrows) cells during alveologenesis (n=4) or homeostasis (n=5), and vehicle (n=3) or naphthalene-injured (n=6) lungs. **i,j,** Quantification of % PDGFRα+ cells with BrdU incorporation. **k,** Freshly sorted cytospin of GFP+ and −/PDGFRα+ cells. l, Histogram of forward scatter (FSC) of PDGFRα+/GFP+ vs. PDGFRα+/GFP− cells (n=16). **m,** Ratio of FSC of GFP+/GFP- cells per lung analyzed. **n,** Percentage of freshly cytospun PDGFRα+ cells with more than one nuclei. **o-q**, Images and quantification of cell size of PDGFRα+ cells grown in culture at passage 0. AW=airway, BM=basement membrane. Scale bars 100um. Each point in graph represents one animal except **q** where each point represent individual cells, with mean ± s.e.m. All p values determined by one-tailed t-test. * p<0.05, ** p<0.01, *** p<0.001, **** p<0.0001.

Next, we set out to define where and when *p16*^*INK4a*^+ cells start to appear in the lung. While senescent cells dependent on *p21*^*CIP1*^ have been described during embryonic development^19,20^, we failed to detect *p16*^*INK4a*^+ cells in the lung during embryogenesis (data not shown). But to our surprise, we detected *p16*^*INK4a*^+ cells in the basement membrane shortly after birth when the oxygen environment dramatically changes in the lung. Thick section images of postnatal day 60 (PND60) lungs showed nuclear GFP staining surrounded by laminin+ basement membrane beneath the airway epithelium (Fig. 1b). We observe these *p16*^*INK4a*^+ cells in similar positions from PND5 during the onset of alveologenesis, to adulthood (defined as >PND45) where the lung is characterized by slow homeostatic cell turnover (Extended Data Fig. 2a-d). The majority of these cells express platelet-derived growth factor receptor alpha (PDGFRα), a fibroblast marker (Fig. 1c,d).

A hallmark of senescence is cell cycle arrest, particularly in cells normally capable of responding to mitogenic stimuli^2^. While this feature is the *sine qua non* of senescence *in vitro*, applying this definition to cells *in vivo* has been difficult due to the lack of effective tools to define and track cells that express biomarkers of senescence in intact tissues. To determine whether *p16*^*INK4a*^+ mesenchymal cells are hypo-replicative *in vivo* relative to *p16*^*INK4a*^-negative mesenchymal cells, we administered continuous BrdU *ad libitum* to separate cohorts of *INKBRITE* animals during the alveologenesis (PND5-45) or adult homeostasis (PND60-150) phases of postnatal lung development and maintenance respectively. Examining PDGFRα+ mesenchymal cells in the sub-airway epithelial compartment, we found a significant reduction of BrdU incorporation in the *p16*^*INK4a*^+ mesenchymal cells during alveologenesis and a trend towards reduction during adult homeostasis compared to *p16*^*INK4a*^-negative mesenchyme (Fig. 1e,f,i, white arrows = PDGFRα+/*p16*^*INK4a*^-/BrdU+, green arrows = PDGFRα+/*p16*^*INK4a*^+/BrdU+). To add a mitogenic stimulus to the lung, which normally exhibit very low cell turnover, we administered naphthalene to injure the lungs of *INKBRITE* animals followed by continuous BrdU administration. Not surprisingly, the airway epithelium demonstrated robust BrdU incorporation during repair (Extended Data Fig. 3a-c), accompanied by an increase in BrdU incoporation in the PDGFRα+ mesenchymal cells (Fig. 1j). However, almost all of the increase in mesenchymal proliferation after naphthalene injury occurred in the *p16*^*INK4a*^-negative mesenchyme, with little to no BrdU incorporation in the *p16*^*INK4a*^+ mesenchyme (Fig. 1g,h,j). To determine whether these cells continue to exhibit cell cycle arrest *ex vivo*, we sorted *p16*^*INK4a*^+ and *p16*^*INK4a*^- PDGFRα+ mesenchymal cells from *INKBRITE* lungs following naphthalene injury and cultured them in proliferative conditions. While *p16*^*INK4a*^-negative mesenchyme continued to proliferate and expand, *p16*^*INK4a*^+ mesenchyme demonstrated significant growth arrest despite exposure to serum-rich media (Extended Fig. 4a-c). These studies demonstrate that *p16*^*INK4a*^+ mesenchymal cells constitute a hypo-replicative population *in vivo* within the basement membrane during postnatal development and homeostasis, and they fail to proliferate upon tissue injury.

Imaging of the *INKBRITE* reporter additionally showed that the nuclear GFP staining often clustered closely together (Extended Fig. 5a), suggesting either *p16*^*INK4a*^+ cells are often adjacent to each other, or a subset of these cells are poly-nucleated *in vivo*. Poly-nucleation is a feature of cellular senescence in cultured fibroblasts^13,21^, but has not been observed in intact tissues. To determine whether poly-nucleated mesenchymal cells exist *in vivo* in the lung, we performed cytospin on PDGFRα+ cells from freshly dissociated adult (>PND60) *INKBRITE* lungs that were either *p16*^*INK4a*^+ or *p16*^*INK4a*^-. Using DAPI and F-actin staining to define the nucleus and cytoplasm respectively, we detected bi- and tri-nucleated mesenchymal cells that are almost exclusively in the *p16*^*INK4a*^+ fraction (Fig. 1k,n). On the cytospin, these cells appeared larger, another common feature of senescent cells^22^. Using forward scatter as an indicator of size, we can demonstrate that freshly sorted *p16*^*INK4a*^+ mesenchyme are significantly larger than their *p16*^*INKa*^- counterpart (Fig. 1l,m). *p16*^*INK4a*^+ mesenchyme maintained these features of senescent cells when cultured *ex vivo*, demonstrating continued enrichment for poly-nucleated cells and enlarged cell size (Fig. 1o, Extended Data Fig. 5b,c). These results demonstrate that *in vivo p16*^*INK4a*^+ mesenchymal cells exhibit features previously ascribed to senescence cells *in vitro*. They also suggest that senescent cells are not absent or rare in healthy and young tissue, but rather present in relative abundance at the epithelial-mesenchymal interface, suggesting a role in maintaining the barrier epithelia in young tissue.

### *p16*^*INK4a*^+ mesenchyme develop a secretory phenotype after epithelial injury

To determine the identity of individual *p16*^*INK4a*^+ cells in the lung and their transcriptional heterogeneity, we performed single cell RNA sequencing (scRNAseq) on sorted GFP+ cells from an adult (PND60) *INKBRITE* lung using droplet-capture approach (10X Chromium Single Cell 3’v2) (Fig. 1a). Cluster analysis of the identified transcripts confirmed our immunohistochemistry, showing that the largest fraction of *p16*^*INK4a*^+ cells is composed of *Pdgfra*+ mesenchymal cells expressing numerous basement membrane components. A smaller fraction is composed of CD45+ immune populations that are predominantly myeloid-derived (mostly alveolar and interstitial macrophages). There is also a very small population of endothelial cells (*Pecam1*+) and negligible epithelial cells (*Epcam*+) (Fig. 2b,c, Extended Data Fig.6).

**Figure 2.**
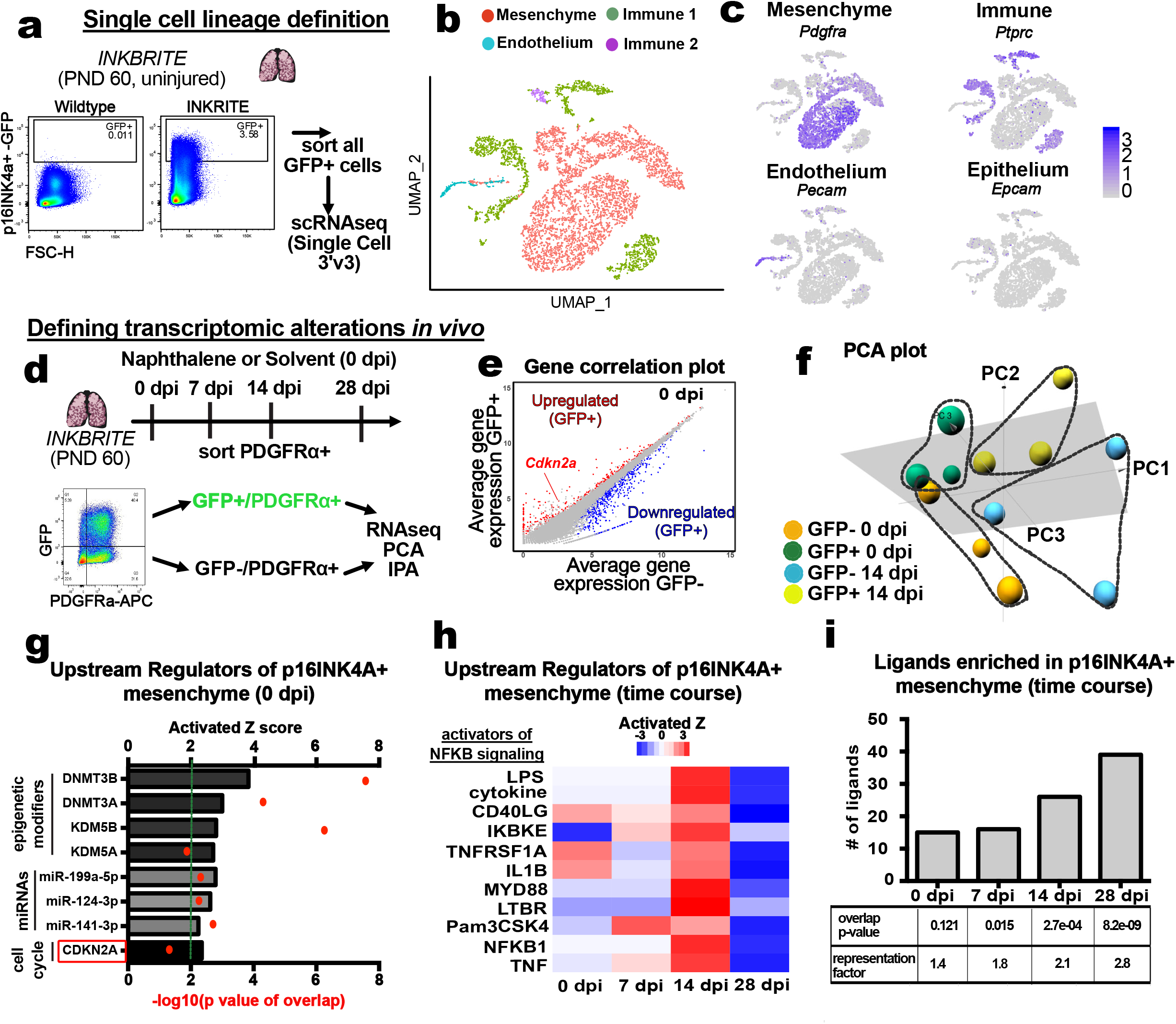
Single cell and bulk RNA sequencing analysis of *p16*^*INK4a*^+ cells in the lung. **a-c,** Single cell analysis of all *p16*^*INK4a*^+ cells in adult uninjured *INKBRITE* lung, showing majority of *p16*^*INK4a*^+ cells are *Pdgfra*+ mesenchymal cells. **d,** Bulk RNAseq of *p16*^*INK4a*^+ and −/PDGFRα+ cells during homeostasis (0 dpi) and injury (7,14,28 dpi) (n=3 per timepoint). **e,** Gene correlation plot showing *Cdkn2a* highly upregulated in GFP+ cells. **f,** PCA plot of *p16*^*INK4a*^+ and – mesenchymal cells at homeostasis and injury. **g,** Predicted upstream regulators driving differentially-expressed genes between *p16*^*INK4a*^+ and – mesenchymal cells during homeostasis. **h,** Heat map of Z scores of upstream regulators activating NF-κB signaling across time points in *p16*^*INK4a*^+ mesenchymal cells. **i,** Number of upregulated genes in *p16*^*INK4a*^+ mesenchymal cells (p <0.05) annotated as a ligand in FANTOM5 database, with corresponding overlap p value and representation factor signifying degree of enrichment.

To more deeply evaluate the transcriptomic signature of *p16*^*INK4a*^+ mesenchyme, we performed bulk RNA sequencing (buRNAseq) on sorted GFP+ or GFP-/PDGFRα+ cells from homeostatic and naphthalene injured (0, 7, 14, and 28 days post injury, or dpi) *INKBRITE* lungs (Fig. 2d). Differentially expressed gene (DEG) analysis (*p16*^*INK4a*^+ vs. *p16*^*INK4a*^-) showed that *Cdkn2a* was highly upregulated at all timepoints in the *p16*^*INK4a*^+ mesenchyme (3’ sequencing can not distinguish between the transcripts of *p16*^*INK4a*^ or *p19*^*ARF*^ encoded by the *Cdkn2a* locus) (Fig. 2e, Extended Data Fig. 7a). Principal component analysis (PCA) revealed clear segregation of identity at the population level between *p16*^*INK4a*^+ and *p16*^*INK4a*^- mesenchyme during homeostasis and injury (Fig. 2f). Ingenuity Pathway Analysis (IPA) of the DEGs for key upstream regulators of the homeostatic (0 dpi) *p16*^*INK4a*^+ cellular transcriptome showed statistically significant activation (Z score > 2) in several epigenetic modifiers and miRNAs that have previously been reported to regulate *p16*^*INK4a*^ expression^23–27^ (*e.g.* KDM5A, KDM5B, DNMT3A, DNMT3B, *etc.*), along with “CDKN2A” itself (Z score = 2.34) as putative regulators of the transcriptional output of *p16*^*INK4a*^+ cells (Fig. 2g).

Time course analysis of the DEGs in *16*^*INK4a*^+ cells demonstrated dynamic changes during the course of repair, notable for a dramatic shift to an inflammatory phenotype after injury. IPA demonstrated a time-dependent activation of upstream regulators associated with NF-κB signaling (*e.g.* LPS, TNFRSF1A, IL1B, MYD88, and *etc.*) specifically in the *16*^*INK4a*^+ mesenchyme that peaks at 14 dpi but dramatically declines at 28 dpi as the injury resolves (Fig. 2h). NF-κB has previously been shown to regulate the senescence-associated secretory phenotype (SASP)^28^, a feature of senescent cells’ ability to secrete soluble ligands. To identify potential paracrine signaling programs induced in *p16*^*INK4a*^+ mesenchymal cells, we cross-referenced DEGs in *p16*^*INK4a*^+ mesenchyme with annotated ligands that have known interacting receptors in the FANTOM5 database^29^. This analysis showed a time-dependent increase in ligands upregulated after injury with significant enrichment at 7, 14, and 28 dpi (Fig. 2i, Extended Data Fig. 7b). These data suggest that *p16*^*INK4a*^+ mesenchyme constitutes a specialized component of the sub-epithelial mesenchyme *in vivo*, with the capacity to respond to inflammatory stimuli during barrier repair by upregulating a secretory program that might modify the regenerative outcome.

### Injured *p16*^*INK4a*^+ mesenchyme enhance epithelial progenitor proliferation *ex vivo*

To determine the effect of *p16*^*INK4a*^+ mesenchyme on the resident stem cells responding to injury, we performed a previously described heterotypic 3D bronchosphere assay^18^ with *p16*^*INK4a*^+ or *p16*^*INK4a*^- mesenchymal (PDGFRα+) cells co-cultured with lineage-labeled *Scgb1a1*+ club cells that proliferate to repair the injured airway epithelium (Fig. 3a). There was no difference in the number of bronchospheres derived from *Scgb1a1*+ club cells when co-cultured with *p16*^*INK4a*^+ or *p16*^*INK4a*^- mesenchyme from homeostatic (0 dpi) *INKBRITE* lungs (Fig. 3b-d). However, *p16*^*INK4a*^+ mesenchyme isolated from naphthalene-injured *INKBRITE* lungs (7 and 14 dpi) demonstrated significantly enhanced capacity to promote bronchosphere formation (Fig. 3e-j). This suggests the presence of SASP-related epithelial mitogen(s) induced upon tissue injury to enhance repair.

**Figure 3.**
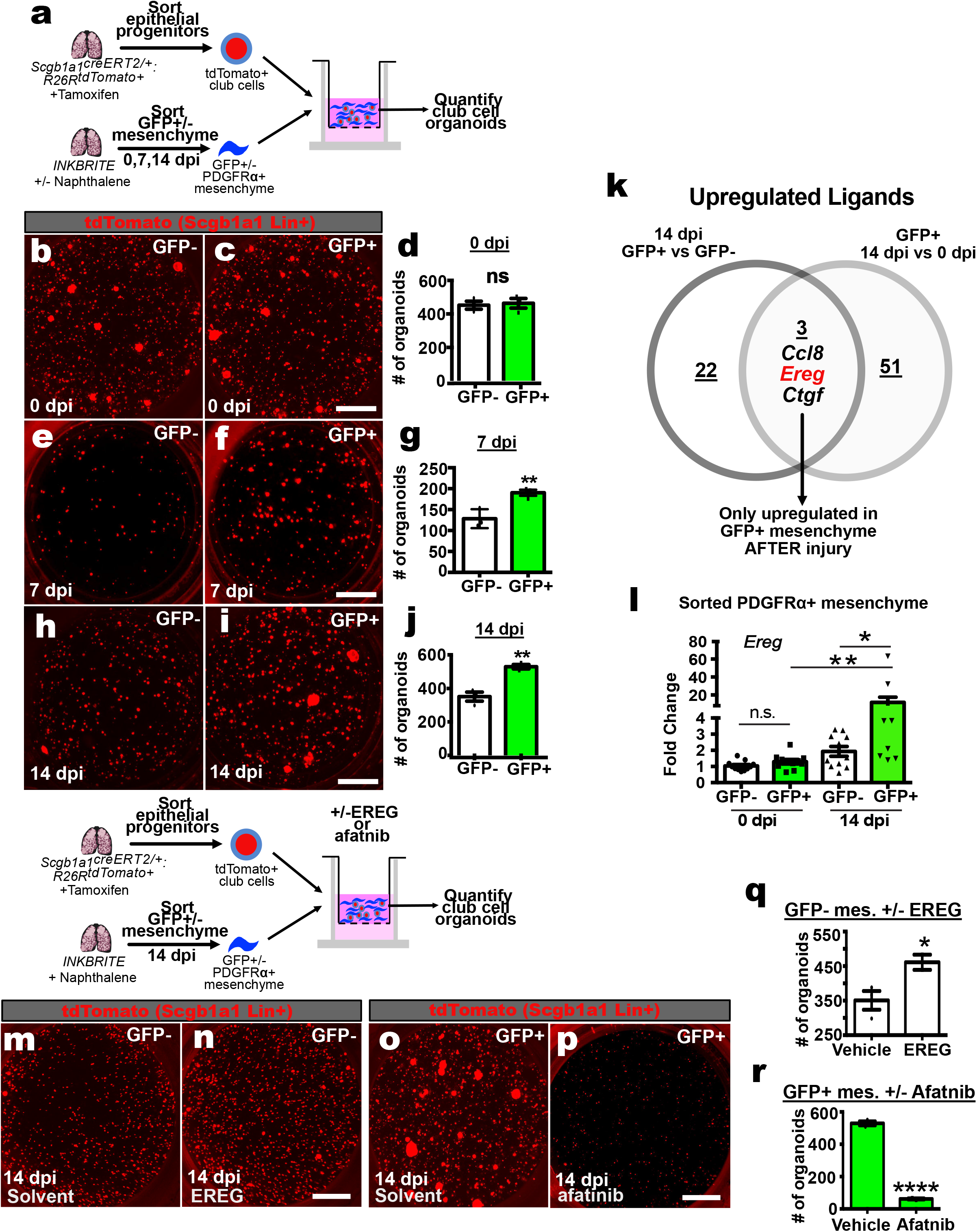
Injured *p16*^*INK4a*^+ mesenchymal cells enhance epithelial progenitor proliferation *ex vivo*. **a,** Heterotypic 3D organoid assay combining uninjured Scgb1a1 Lin+ epithelial progenitors with *p16*^*INK4a*^+ or −/PDGFRα+ cells during homeostasis or injury. **b-j,** Images and quantification of Scgb1a1 Lin+ organoid growth co-cultured with *p16*^*INK4a*^+ or – mesenchymal cells during homeostasis (0 dpi) or injury (7,14 dpi). **k,** Identification of FANTOM5 annotated ligands upregulated only in GFP+ mesenchyme after injury. **l,** qPCR of *Ereg* on sorted GFP+ or −/ PDGFRα+ mesenchyme from *INKBRITE* lungs at homeostasis or injury. **m,n,** Images of Scgb1a1 Lin+ organoids co-cultured with injured (14dpi) GFP- mesenchyme +/− recombinant EREG. **o,p,** Images of Scgb1a1 Lin+ organoids co-cultured with injured (14dpi) GFP+ mesenchyme +/− Afatnib. **q,r,** Quantification of organoids from **m-p**. Scale bars 100um. Each point in graph represents one well with mean ± s.e.m. Experiments performed in triplicates. All p values determined by one-tailed t-test. * p<0.05, ** p<0.01.

To determine paracrine factors that might be activated specifically in *p16*^*INK4a*^+ mesenchyme only after injury, we analyzed DEGs that are upregulated (p <0.05) in the *p16*^*INK4a*^+ mesenchyme at 14 dpi when compared to both *p16*^*INK4a*^- mesenchyme at 14 dpi and *p16*^*INK4a*^+ mesenchyme at homeostasis (0 dpi). This overlapping gene set represents genes that are preferentially upregulated in *p16*^*INK4a*^+ mesenchyme only after injury, and cross-referencing with annotated ligands from FANTOM5 revealed 3 ligands, with one of them (*Ereg*) encoding a protein named epiregulin (Fig. 3k, Extended Data Fig. 7a). qPCR analysis of sorted mesenchyme at 0 and 14 dpi confirmed that *Ereg* is upregulated only in the *p16*^*INK4a*^+ mesenchyme after injury (Fig. 3l). EREG signals through the receptors EGFR and ERBB4, and is reported to be a potent hepatocyte and keratinocyte mitogen^30–32^, but its source and role in the lung is unclear. We added recombinant EREG to injured (14 dpi) *p16*^*INK4a*^-negative mesenchyme co-cultured with *Scgb1a1*+ club cells, which significantly enhanced bronchosphere growth (Fig. 3m,n,q). Conversely, afatnib, a small-molecule antagonist of EGFR and ERBB4, dramatically attenuated bronchosphere growth when added to the organoid culture with injured (14 dpi) *p16*^*INK4a*^+ mesenchyme (Fig. 3o,p,r). These experiments demonstrate that *p16*^*INK4a*^+ cells in the sub-epithelial mesenchyme stands poised to promote epithelial proliferation after injury by secreting a novel club cell mitogen, EREG, to enhance airway barrier repair.

### Mesenchymal *p16*^*INK4a*^ is required for epithelial regeneration in vivo

IPA analysis of *p16*^*INK4a*^+ mesenchyme shows *Cdkn2a* as a major upstream regulator (Fig. 2g), suggesting that the transcription of *p16*^*INK4a*^ from this locus accounts for its unique function, including the secretory program associated with injury. To determine the specific role of *p16*^*INK4a*^ in modifying mesenchymal feedback to the epithelium, we deleted *p16*^*INK4a*^ using a mesenchymal-specific Cre-driver (*Dermo1*^*Cre/+*^)^33^. Mesenchymal-specific deletion of *p16*^*INK4a*^ (*Dermo1*^*Cre/+*^:*p16*^*flox/flox*^, referred to as *Dermo1*^*p16CKO*^) did not alter the gross morphology of the adult lung (Fig. 4a,b,e,f), suggesting it is not necessary for postnatal development. However, after injury with naphthalene, *Dermo1*^*p16CKO*^ airways demonstrated a dramatic reduction in epithelial repair as quantified by the number of SCGB1A1+ club cells present as well as qPCR of *Scgb1a1* transcripts in the whole lung (Fig. 4c,d,e,f), leaving β4-tubulin+ ciliated cells as the remaining major airway epithelial population.

**Figure 4.**
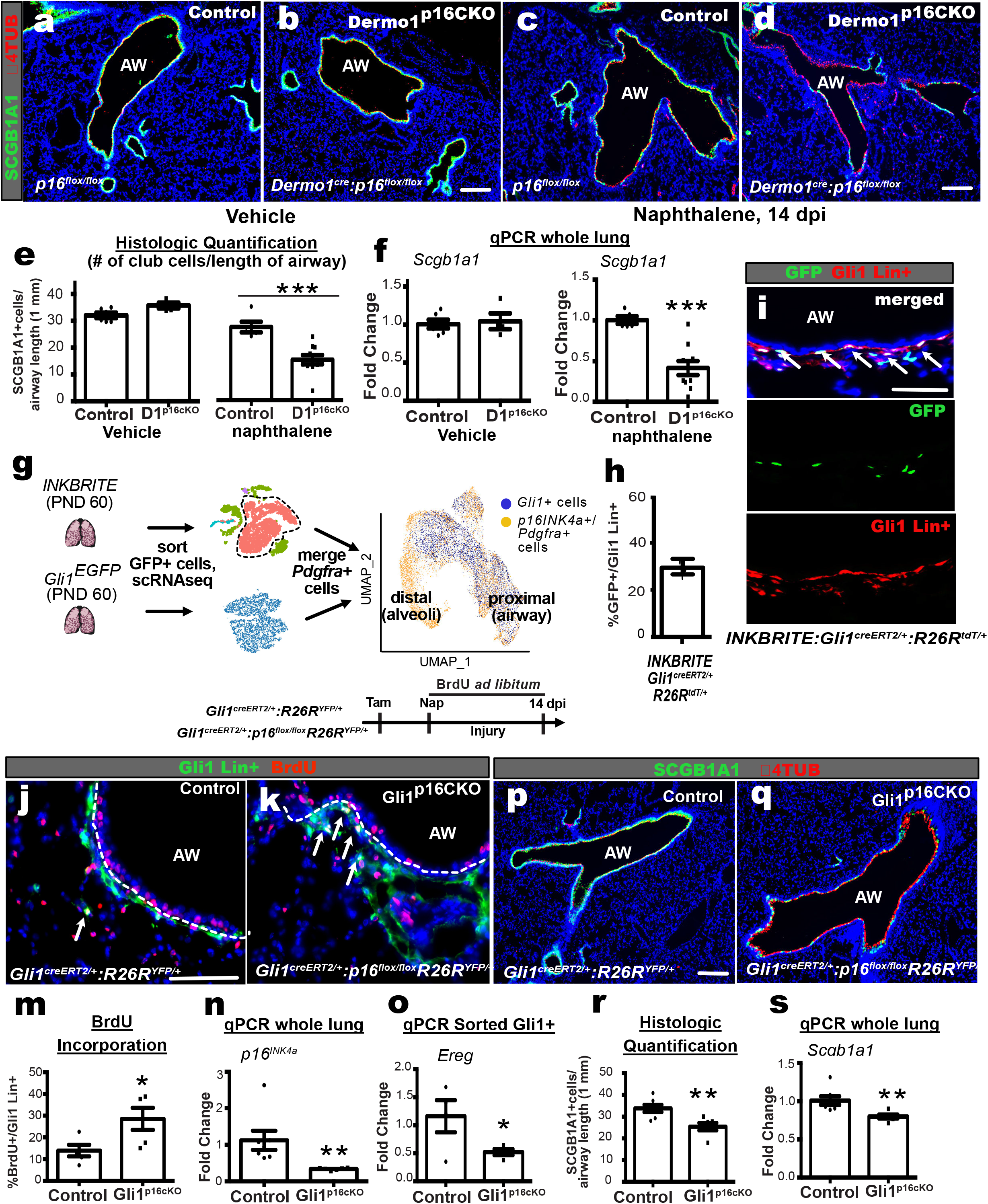
Mesenchymal *p16*^*INK4a*^ is required for epithelial regeneration *in vivo*. **a-d,** Images of the airways of control (n=7) and *Dermo1*^*p16CKO*^ (n=9) lungs with and without naphthalene injury. **e,** Histological quantification of SCGB1A1 club cells per unit of airway length (1mm). **f,** qPCR of *Scgb1a1* of whole lung RNA. **g,** Merge of *p16*^*INK4a*^+/*Pdgfra*+ single cell transcriptome with Gli1+ single cell transcriptome and projected on UMAP. **i,h,** Co-localization of *Gli1* Lin+ cells with GFP in *INKBRITE:Gli1*^*creERT2*^*:R26R*^*tdT/+*^ lungs and quantification of the overlap. (n=3). **j-m,** Images and quantification of BrdU incorporation in Gli1 Lin+ cells (white arrows) of control (n=5) and *Gli1*^*p16CKO*^ lungs (n=5) after naphthalene injury. **n,** qPCR of *p16*^*INK4a*^ in whole lung RNA. **o,** qPCR of *Ereg* of sorted Gli1 Lin+ cells after naphthalene. **p-s,** Images, histological quantification of SCGB1A1 club cells, and qPCR of *Scgb1a1* on control (n=7) and *Gli1*^*p16CKO*^ (n=6) lungs after naphthalene injury. AW=airway. Scale bars 100um. Each point in graph represents one animal with mean ± s.e.m. All p values determined by one-tailed t-test. * p<0.05, ** p<0.01, *** p<0.001.

The *Dermo1* Cre-driver is active in a broad and heterogeneous mesenchymal population. To further refine the *p16*^*INK4a*^+ population that might influence airway progenitor behavior, we compared *p16*^*INK4a*^+ mesenchyme scRNAseq data with those captured from the lung of a *Gli1* reporter (*Gli1*^*EGFP*^). *Gli1* marks a subset of cells possessing mesenchymal stromal cell properties *ex vivo^34^,* and capacity to modulate neighboring epithelium and immune cells *in vivo*^18,35^. Merge of *p16*^*INK4a*^+/*Pdgfra*+ cells with *Gli1*+ cells followed by unsupervised clustering shows that *Gli1*+ cells comprise a subset of *p16*^*INK4a*^+ mesenchyme enriched for previously reported proximal airway mesenchymal markers^36^ (Fig. 4g, Extended Data Fig. 8). To confirm co-localization of *Gli1* and *p16*^*INK4a*^, we generated *INKBRITE*:*Gli1*^*CreERT2/+*^:*R26R*^*Tdt/+*^ triple transgenic animals which shows a high degree of overlap of the two markers around the airway epithelium (Fig. 4h,I, ~30% of *Gli1* Lin+ also GFP+).

We then generated *Gli1*^*CreERT2/+*^:*p16*^*flox/flox*^:*R26R*^*YFP/+*^ (referred to as *Gli1*^*p16CKO*^) animals to delete *p16*^*INK4a*^ in a more temporal and spatial-specific manner and track the behavior of *p16*^*INK4a*^ deleted-lineage labeled *Gli1*+ (*Gli1* Lin+) mesenchyme. Tamoxifen induction in young (2 months old) *Gli1*^*p16CKO*^ animals caused a significant reduction of *p16*^*INK4a*^ transcripts compared to controls (*Gli1*^*CreERT2/+*^:*R26R*^*YFP/+*^) (Fig. 4n). Continuous BrdU administration to *Gli1*^*p16CKO*^ animals after naphthalene injury showed that *p16*^*INK4a*^ deletion increased the proliferation of *Gli1* Lin+ cells following injury (Fig. 4j,k, m), but reduced the expression of the epithelial mitogen *Ereg* (Fig. 4o). Similar to the *Dermo1*^*p16CKO*^ animals, *Gli1*^*p16CKO*^ animals experienced reduced airway epithelial progenitor recovery after injury (Fig. 4p-s). These data suggest that *p16*^*INKa*^ is required to maintain cell-cycle arrest and secretory function in senescent mesenchymal cells *in vivo*, the loss of which disrupts mesenchymal signals to progenitors that ensure repair of the barrier epithelia.

## Discussion

We generated an ultra-sensitive reporter mouse that allows us to isolate and localize *p16*^*INK4a*^+ cells in tissues. While there are numerous biomarkers for cellular senescence, we focused on *p16*^*INK4a*^ because of the reported benefits of eliminating *p16*^*INK4a*^+ cells from aging tissues. This has led some to label these cells as “zombie cells^37^,” a popular term to describe senescent cells in the lay press. However, our data suggest that the term “zombie cell” may be inaccurate, as we describe cells with senescent features that remain responsive to physiological cues to maintain organ integrity. Leveraging our novel reporter, our data show that *p16*^*INK4a*^+ cells arise normally during postnatal tissue maturation in mesenchyme producing basement membrane components, and function as sentinels within the stem cell niche that activate upon barrier injury to promote epithelial repair (Extended Data Fig. 9). Contrary to the prevailing idea that *p16*^*INK4a*^ transcript is absent in young and healthy tissues, our data suggest that *p16*^*INK4a*^ is continuously present throughout the lifespan (albeit at a very low level) to maintain senescence in a mesenchymal subset poised to respond to epithelial injury. Furthermore, our work shows that senescent characteristics can be acquired in stages *in vivo*, as we demonstrated that *p16*^*INK4a*^+ cells acquire SASP-like behavior only after tissue damage in young organs. This suggests that the term “presenescent,” previously used to describe early passage non-senescent fibroblasts in culture^38^, maybe better applied to hypo-replicative fibroblasts *in vivo* that are poised to transition to a secretory fate. Our ability to isolate and study *p16*^*INK4a*^+ cells at the cellular resolution *in vivo* provides a framework to better understand the physiological role of senescence across tissues and life span. While further work is needed to determine how these *p16*^*INK4a*^+ cells change during organ aging, and how that modifies their sensitivity to different classes of senolytics, our work suggests a potential cost to the elimination of *p16*^*INK4a*^+ cells that needs to be considered in therapeutic trials.

---

**Extended Data Figure 1.**
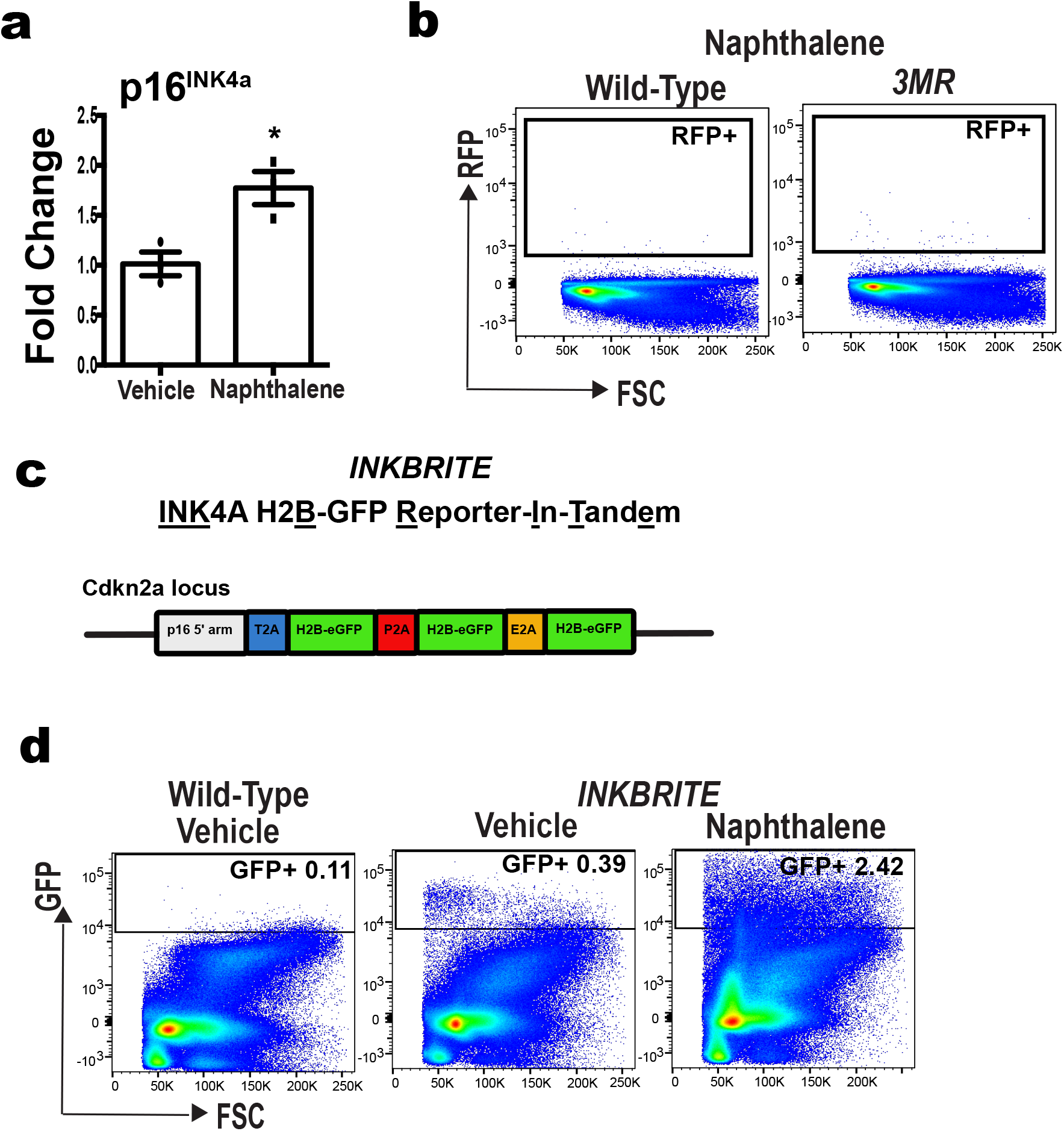
Comparison of *3MR* and *INKBRITE* reporters. **a,** qPCR of *p16*^*INK4a*^ in whole lung RNA of vehicle (n=3) vs. naphthalene (n=3) treated wild type lungs at 14dpi. **b,** FACS analysis of *3MR* lungs after naphthalene injury. **c,** Target construct design for *INKBRITE*. **d,** FACS analysis of wild type and *INKBRITE* lungs before and after naphthalene injury. Each point in graph represents one animal with mean ± s.e.m. All p values determined by one-tailed t-test. * p<0.05.

**Extended Data Figure 2.**
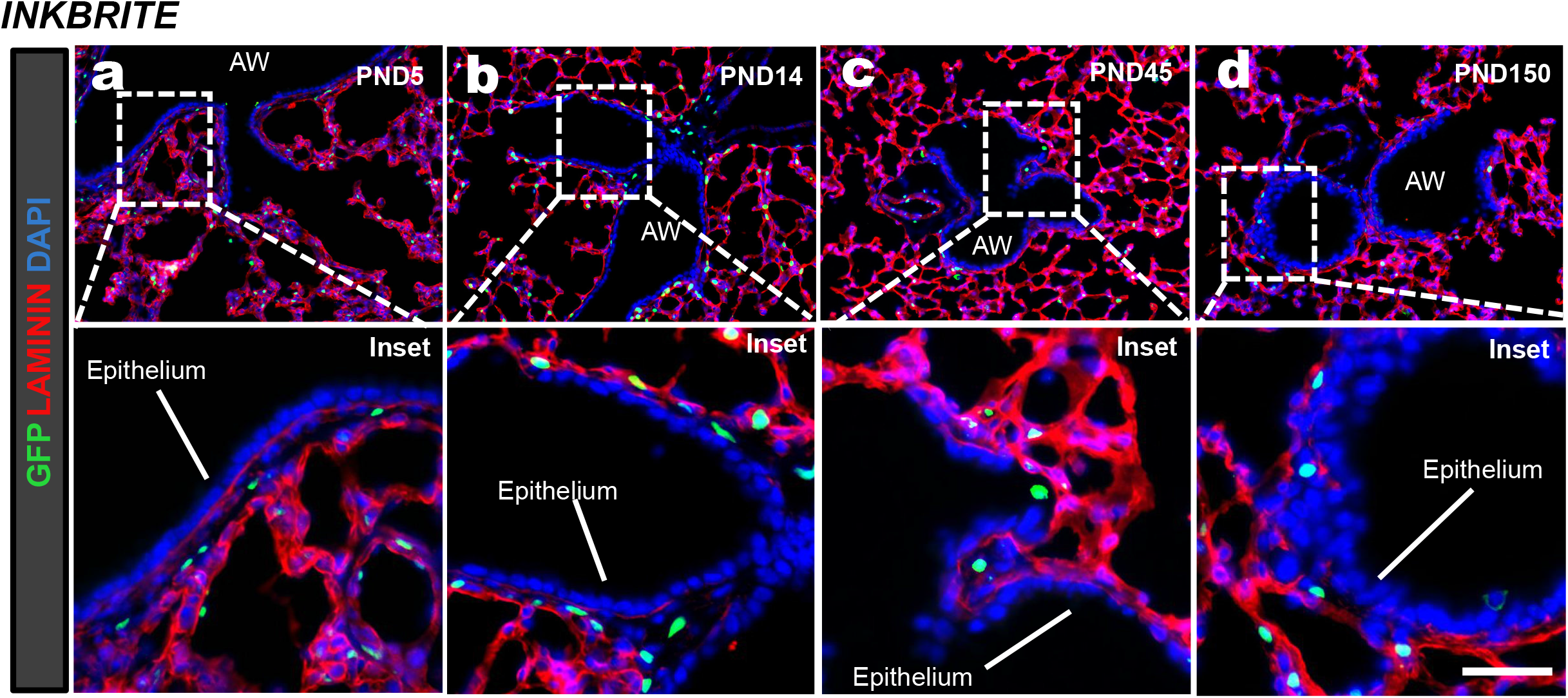
Localization of GFP+ cells in *INKBRITE* lungs across time. **a-d,** IHC of GFP overlaid with laminin during alveologenesis (PND5,14) and homeostasis (PND45,150) phases of postnatal lung maturation. AW=airway, Scale bars 100um.

**Extended Data Figure 3.**
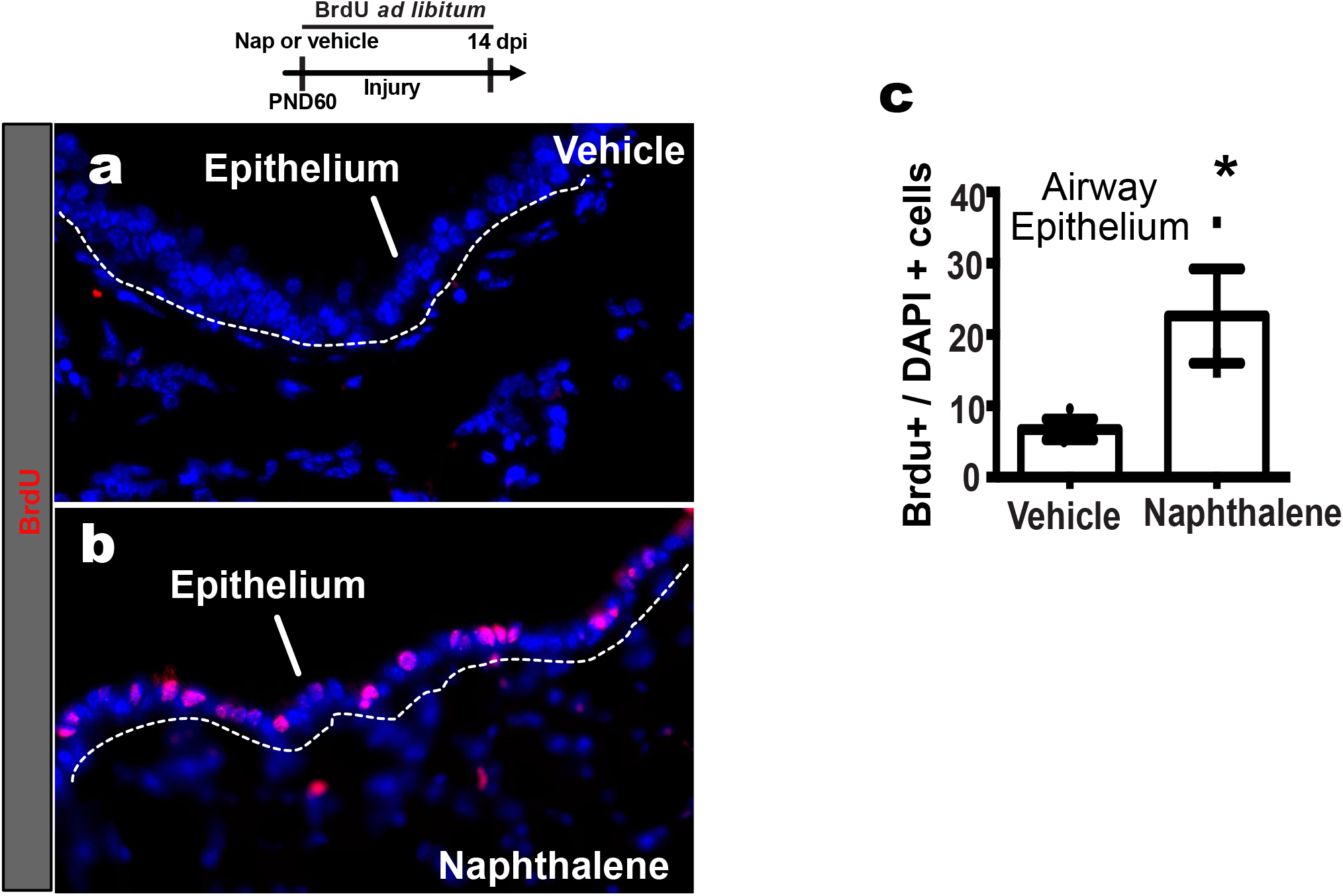
Airway epithelial proliferation after injury. **a-c,** IHC and quantification of BrdU incorporated into airway epithelium with vehicle (n=3) or naphthalene-injured lungs (n=3). AW=airway. Scale bars 100um. Each point in graph represents one animal with mean ± s.e.m. All p values determined by one-tailed t-test. * p<0.05.

**Extended Data Figure 4.**
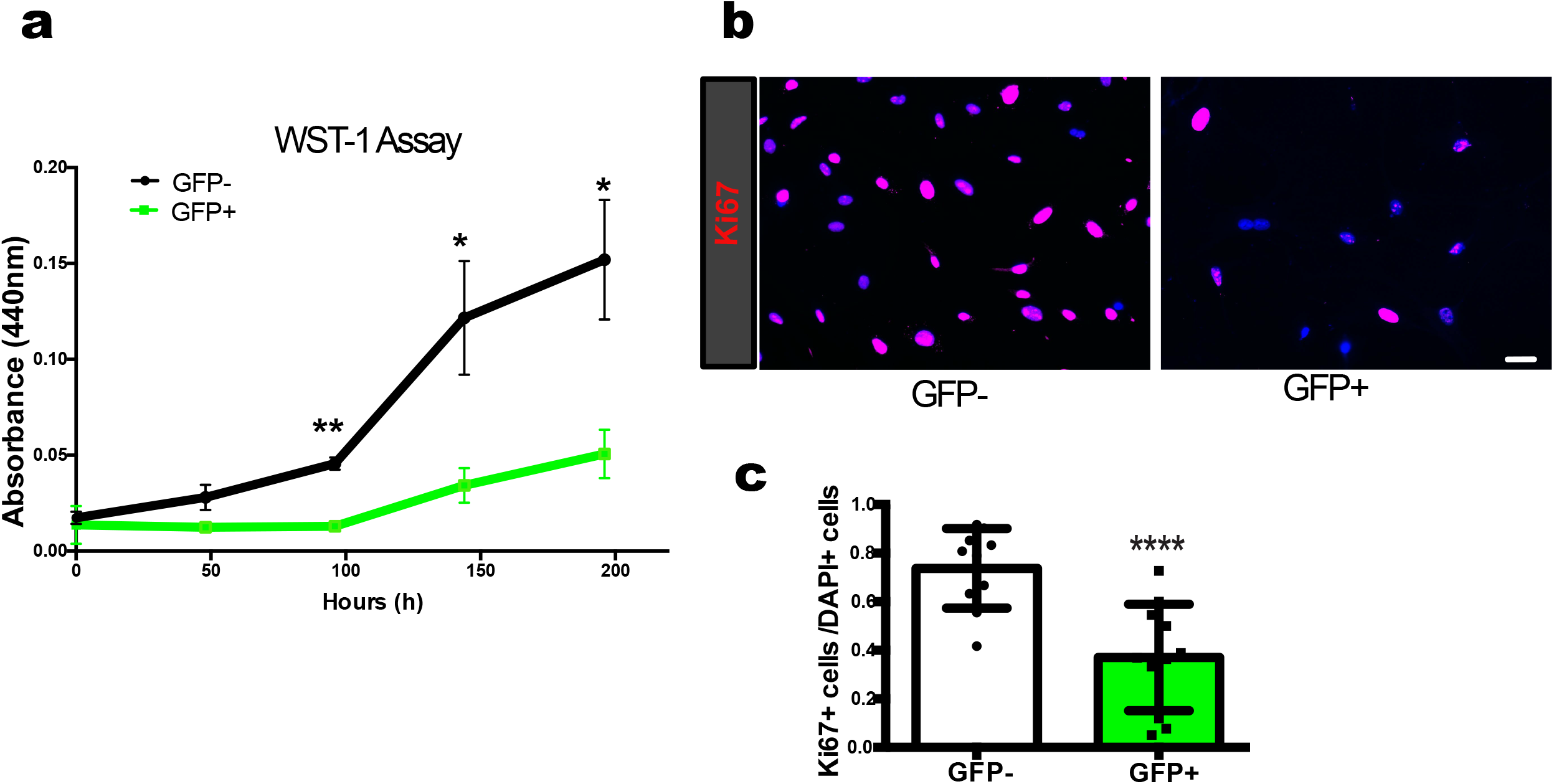
Proliferation of GFP+ mesenchymal cells sorted from *INKBRITE* lungs after injury. **a,** WST-1 proliferation assays on sorted GFP+ or −/PDGFRα+ mesenchyme from *INKBRITE* lungs after naphthalene injury. **b,c,** Images and quantification of Ki67 on sorted GFP+ or −/PDGFRα+ mesenchyme from *INKBRITE* lungs. Scale bars 100um. Each point in graph represents one well with mean ± s.e.m. All p values determined by one-tailed t-test. * p<0.05, ** p<0.01, **** p<0.0001.

**Extended Data Figure 5.**
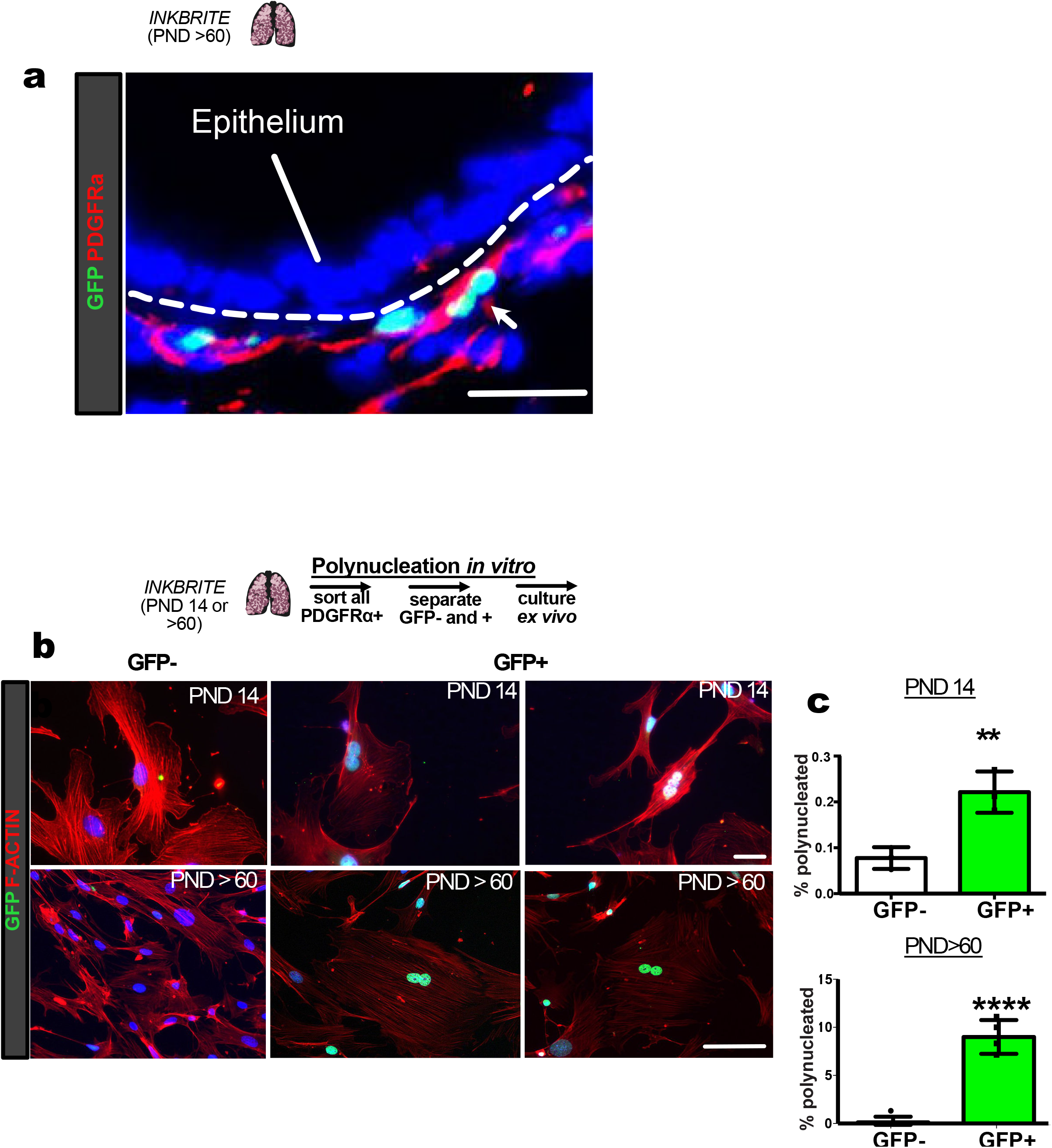
Identification of poly-nucleated *p16*^*INK4a*^+ cells *in vivo* and *in vitro*. **a,** Image of closely adjoined nuclei of GFP+/PDGFRα+ cell or cells in the uninjured *INKBRITE* lung. **b,c,** Images and quantification of nuclei on sorted GFP+ or −/PDGFRα+ mesenchyme from *INKBRITE* at PND14 and >60 lungs. Scale bars 100um. Each point in graph represents one well with mean ± s.e.m. All p values determined by one-tailed t-test. ** p<0.01, **** p<0.0001.

**Extended Data Figure 6.**
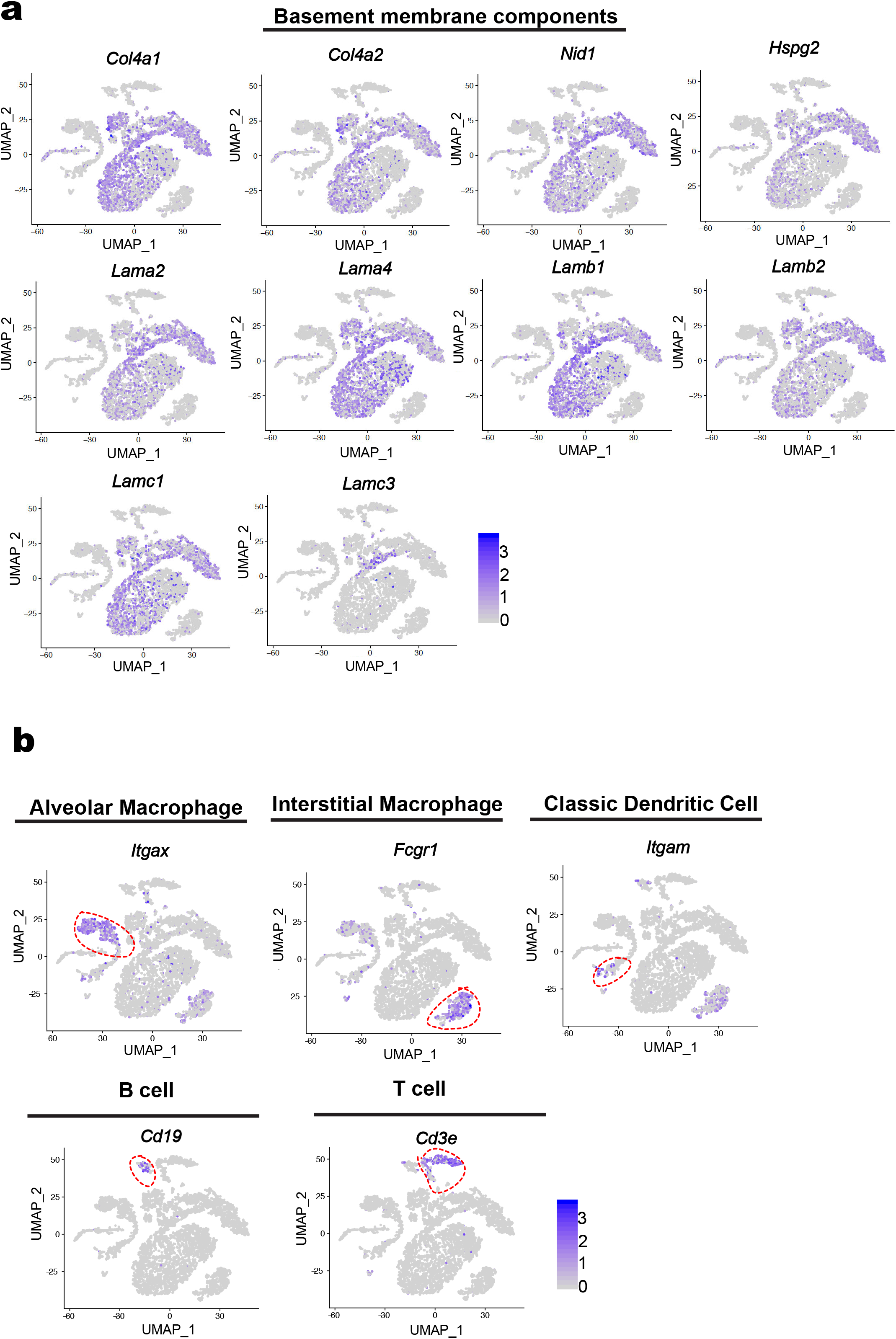
Feature plots of gene expression in *p16*^*INK4a*^+ cells in the lung. **a,** Expression of basement membrane components in the *Pdgfra*+ cluster of *p16*^*INK4a*^+ cells. **b,** Myeloid and lymphoid lineages of *p16*^*INK4a*^+ immune cells.

**Extended Data Figure 7.**
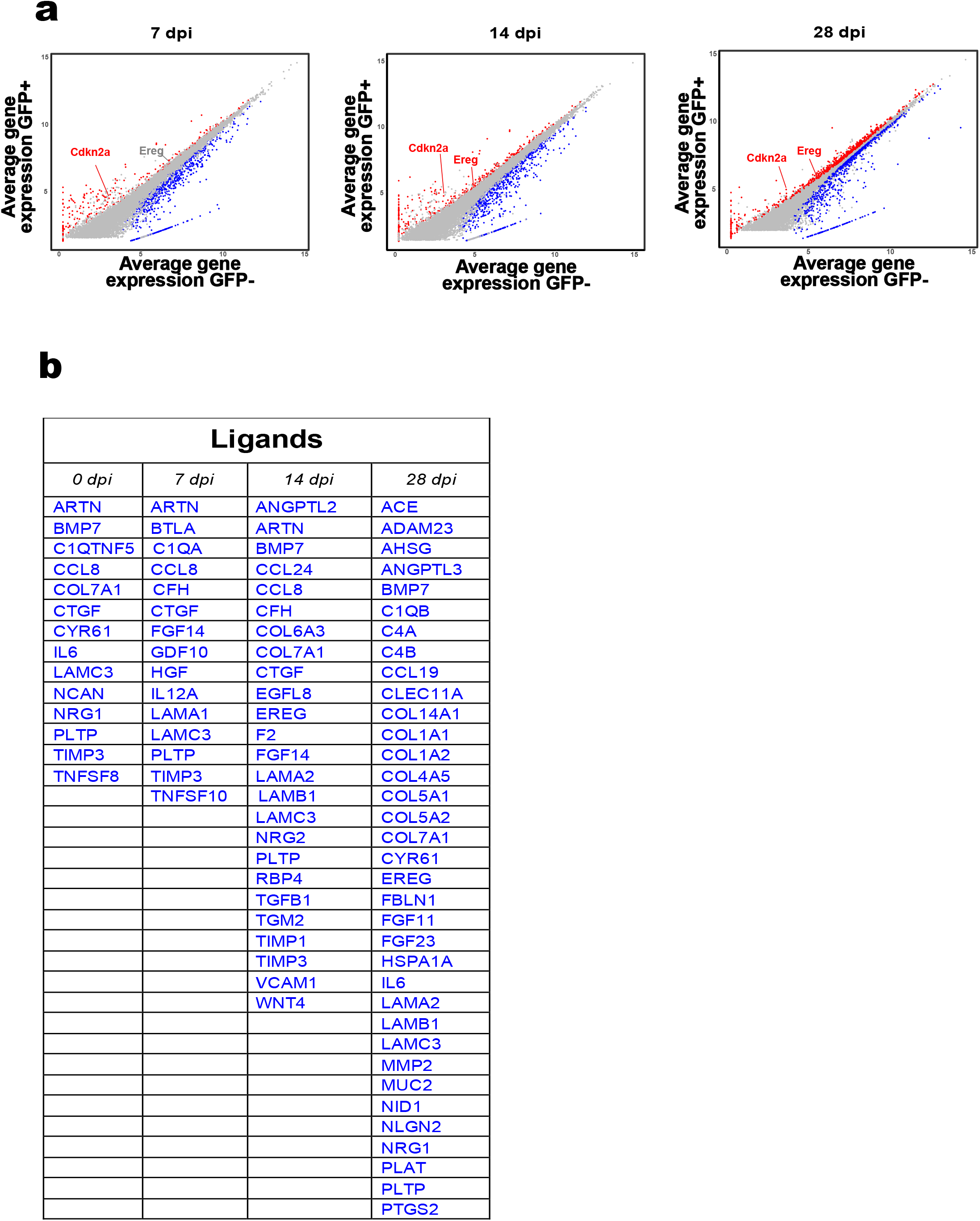
Bulk RNAseq analysis of *p16*^*INK4a*^+ mesenchymal cells in the lung. **a,** Gene correlation plot showing expression of *Cdkn2a* and *Ereg* after injury. **b,** Table of FANTOM5 annotated ligands upregulated in *p16*^*INK4a*^+ mesenchymal cells during homeostasis and injury.

**Extended Data Figure 8.**
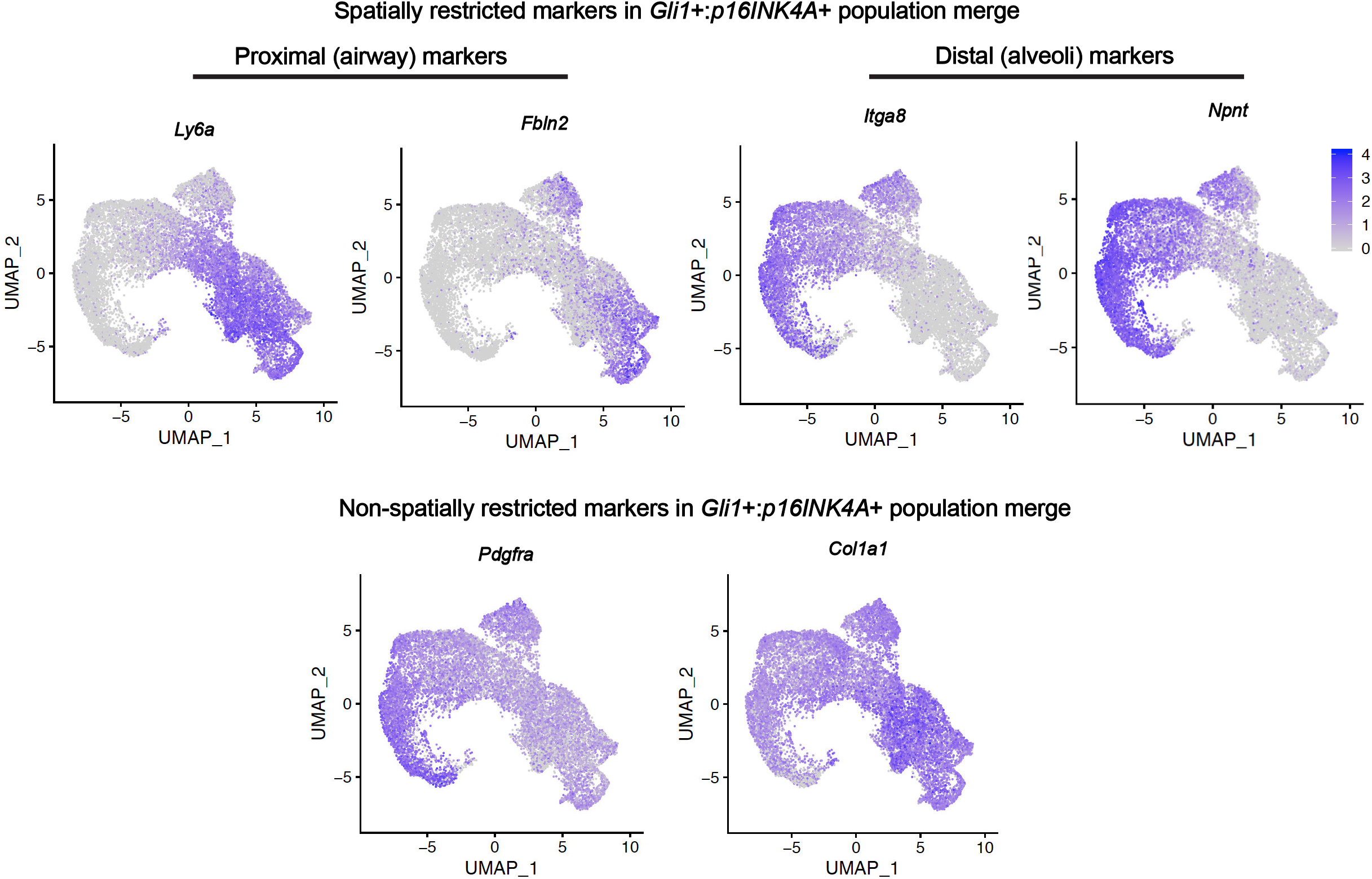
Feature plots of gene expression in *Gli1+*/ *p16*^*INK4a*^+ population merge in the lung.

**Extended Data Figure 9.**
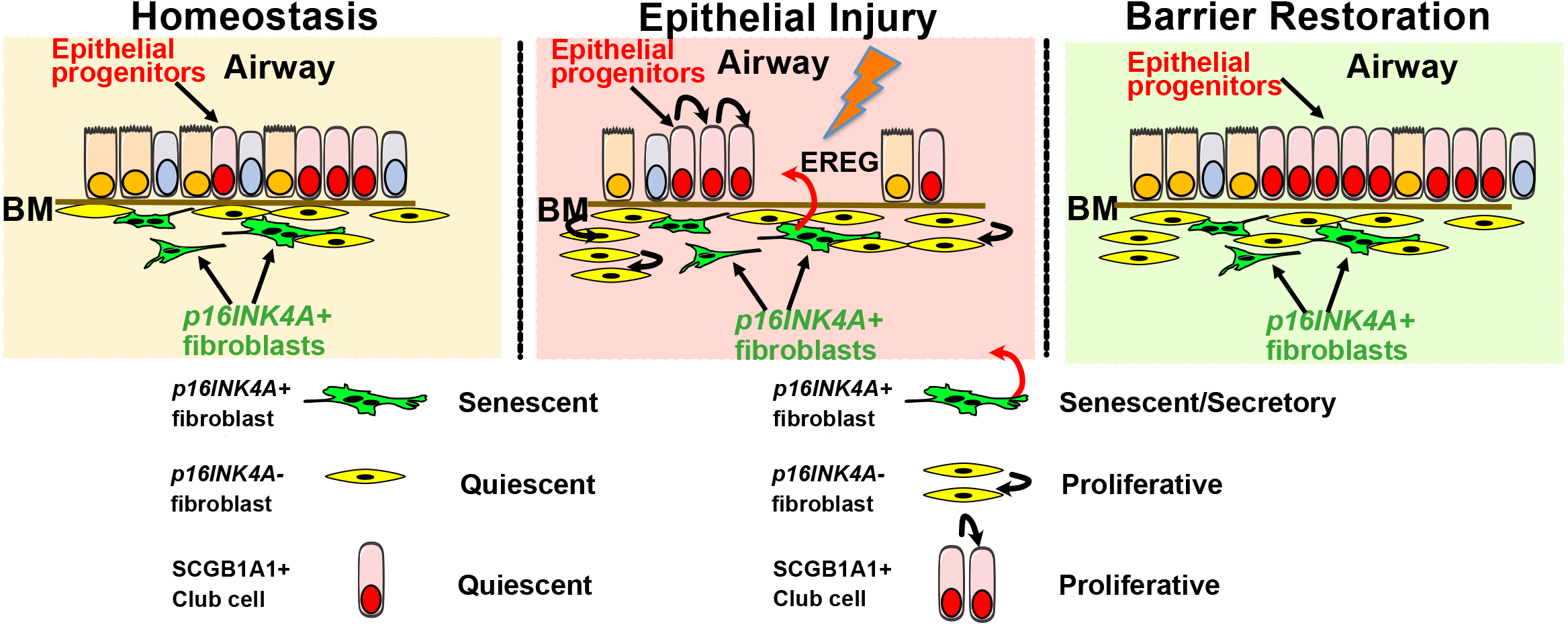
Model of *p16*^*INK4a*^+ mesenchymal cells acting as sentinels poised to enhance epithelial repair after barrier injury.

## Methods

### Generation of INKBRITE mouse model

Three H2B-GFP cassettes were cloned in tandem into one cassette with each separated by a 2A self-cleaving peptide (3X 2A-H2B-GFP). The 3X-2A-H2B-GFP fusion cassette was then inserted into bacterial artificial chromosome (BAC) containing the *Cdkn2a* locus (BAC clone RP24-322D20 from the RPCI-24 C57BL/6J male mus musculus BAC library http://bacpac.chori.org/library.php). The 3X 2A-H2B-GFP cassette was inserted in frame with the p16INK4A reading frame in exon 2 at the identical insertion site as the 3MR model as previously described^1^. The insertion site prevents the production of a full length p16INK4a transcripts and also produces a premature stop codon in the p19ARF reading frame within the shared exon 2 of the two transcripts, so no mature transcripts of p16 or p19 are produced by the BAC. Furthermore, the insertion frame of the cassette would produce a frame shift mutation in the p19ARF reading frame without translation of the H2B-GFP protein, thus only p16INK4A expression is reported. A neo cassette 3’ to the 3X 2A-H2B-GFP cassette used for selection was removed before the BAC injection. Pronuclear injection of the BAC was performed in eggs from C57bl/6 strain donors and implanted to establish founders. One single founder sequenced to determine a single integration site in chromosome 18 was used for all subsequent experiments.

### Animal Studies

All animals were housed and treated in accordance with the Institutional Animal Care and Use Committee (IACUC) protocol approved at the University of California, San Francisco Laboratory Animal Resource Center (LARC). Generation and genotyping of the *Gli1*^*cre/ERT2*^, *R26R*^*EYFP*^, *R26R*^*tdTomato*^, *Scgb1a1*^*creERT2*^, Dermo1^cre^ lines were performed as previously described by The Jackson Laboratory. The *p16*^*flox/flox*^ allele as previously described was a gift from Ned Sharpless. 8-12 week old littermate mice were gender balanced and randomly assigned to experimental groups. A minimum of three mice was used in each experimental group.

For lineage tracing studies, tamoxifen was dissolved in corn oil and administered intraperitoneally (IP) at 200 mg/kg body weight per day for five consecutive days. Animals were injured with naphthalene by administering naphthalene dissolved in corn oil IP at 150-300 mg/kg one time. Lung tissue was collected at 7, 14, or 28 days post injury. BrdU administration was performed *ad libitum* at a concentration of 0.5 mg/ml with 1% sucrose in dH_2_O. Brdu water was refreshed twice a week for the duration of each experimental period.

### Histology and Immunofluorescence

Detailed description on harvesting the lung, cryo/paraffin embedding of tissues, sectioning, and immunohistochemistry methods are previously described^2^. Primary antibodies and dilution used for histology include: anti-GFP 1:250 (chicken, Aves Labs, #1020, and goat, Abcam, #6673), anti-SCGB1A1 1:500 (goat, Santa Cruz, #9772), anti-beta IV Tubulin 1:200 (mouse, Abcam, #11315), anti-laminin 1:200 (rabbit, Sigma, #9932), anti-PDGFRα (rabbit, Cell Signaling, #3174), anti-BrdU 1:200 (rat, Abcam, #6326). Sections were imaged for quantification on a Zeiss Lumar V12 microscope. At least three samples per genotype/condition were used, and at least 5-10 randomly selected sections were chosen for each sample. Cell counts for SGCGB1A1+, Brdu+, GFP+, tdTomato+, and DAPI cells were performed on Fiji using the “Cell Counter” plug-in. For quantification of airway length, the freehand line feature coupled with “Measure” was utilized. Images were blinded to experimental condition for quantification.

### Cleared Thick Slice Imaging

Mouse lung was extracted as above and fixed in 4% PFA overnight at 4 °C. Tissue was washed with changes of 1X PBS for 2 hours and cut into 200 μm sections on a vibratome. Sections were blocked for at least 1 hour in 0.3% Triton X, 5% FBS, 0.5% BSA in PBS with azide at 4 °C. Sections were then incubated in primary antibodies (Key Resources Table) in PBS with 0.15% Triton X, 7.5% FBS, and 0.75% BSA for 1-3 days at 4 °C. Sections were washed with 0.15% Triton X and incubated in secondary antibodies (Key Resources Table) overnight at 4 °C. After several washes with 0.15% Triton X, DAPI was added in PBS for 30 minutes and sections were rinsed again. Sections were cleared using RIMS (40 g Histodenz, 30 mL PBS, 5 μL Tween 20, 50 μL 10% azide) for 30-60 min and mounted. Images were taken on a Nikon A1R multi-photon confocal microscope. Surfaces were created for each marker of interest to generate images.

### Lung digestion and Fluorescence Activated Cell Sorting (FACS)

For mouse, whole lung was dissected from adult animals and tracheally perfused with a digestion cocktail of Collagenase Type I (225 U/mL), Dispase (15 U/mL) and Dnase (50 U/mL) and removed from the chest. The lung was further diced with a razor blade and incubated in digestion cocktail for 45 mins at 37 °C with continuous shaking. The mixture was then washed with sorting buffer (2% FBS and 1% Penicillin-Streptomycin in DMEM). The mixture was passed through a 70 μm cell strainer and resuspended in RBC lysis buffer, then passed through a 40 μm cell strainer. Cell suspensions were incubated with the appropriate conjugated antibodies in sorting buffer for 30 min at 4 °C and washed with sorting buffer. Doublets and dead cells were excluded based on forward and side scatter and DRAQ7 (Cat#7406S; Cell Signaling Technologies) fluorescence, respectively. Immune and endothelial cells were excluded using anti-CD45 (Alexa Flour 700; Cat#560510; BD; used 1:200) and anti-CD31 (APC/Fire750; Cat#102528; BioLegend; used 1:200), respectively. Epithelial cells were also excluded using anti-CD326/Epcam (BV421; Cat#563214; BD; used 1:200), and finally, PDGFRα+ cells were included (APC; Cat#17-1401-81; Thermo; used 1:200). After selection for PDGFRα+ cells, the GFP- and GFP+ mesenchyme were further separated and collected into sorting buffer. For *Scgb1a1*^*creERT2*^*:R*^*tdt*^ epithelial cells, a live/dead stain was done first then tdTomato+ cells were sorted and collected into sorting buffer. Analysis was performed using FlowJo software.

### Single Cell Sequencing and Analysis

Single cell suspension of the lung was prepped and sorted as described above. Cells were loaded onto a single lane per sample as 1000cells/μl into the Chromium™ Controller to produce gel bead-in emulsions (GEMs). GEMs underwent reverse transcription for RNA barcoding and cDNA amplification, with the library prepped using the Chromium Single Cell 3’ Reagent Version 3 kit. Each sample was sequenced in 1 lane of the HiSeq2500 (Illumina) in Rapid Run Mode. To build transcript profiles of individual cells the CellRanger v3.0.3 software with default settings was used for de-multiplexing, aligning reads with STAR software to mouse genome GRCm38, and counting unique molecular identifiers (UMIs). We used the Seurat R package along with a gene-barcode matrix provided by CellRanger for downstream analysis. In total, we filtered the data in 2 different steps. We first filtered the dataset by only accepting cells that expressed a minimum of 200 genes and genes that were expressed in at least 3 cells. The UMI were log-normalized and we to the identified genes with high expression and those are variable we used the mean variance relationship method. Our second filtered was set to accept cells with less than 6500 unique gene counts and 15% mitochondrial gene counts. Using regress out function, we mitigate the effect of mitochondrial gene counts. Next we used principle component analysis (PCA) to identify components that can be found within our dataset for unsupervised clustering. We used the JackStrawPlot function in the Seurat function to create Scree plots were and compare p-value (significance) for each PC. We selected 10 different PCA’s for clustering of the 4600 cells. Clustering results were visualize using the Uniform Manifold Approximation and Projection (UMAP) algorithm in the Seurat package. 4 clusters were identified in p16+ cells. We further integrated the PDGFRA+ clusters from both p16-GFP+ and GLI1+ cells scRNAseq data sets by using IntegrateData function in Seurat v3.0. We performed an integrated analysis on all cells. The standard workflow was carried out using ScaleData, RunPCA, RunUMAP, FindNeighbors and FindClusters functions to cluster cells. For individual gene visualization on all clusters we used the FeaturePlot function.

### Bulk RNA Sequencing and Analysis

For bulk RNAseq sequencing was done using Sanger/Illumina 1.9, and there were an average of 45 million reads per sample with a total of 3 biological replicates per condition. Quality control of reads was conducted by using FastQC (Babraham Bioinformatics). Ligation adaptors were removed using the Cutadapt and Sickle. Sequencing reads were aligned using HISAT and assembled with Stringtie software to the reference genome *Mus musculus*, UCSC version mm10. All gene counts of the biological replicates were concatenated while running DEseq2 for differential gene expression (DGE). Upstream regulators were generated through the use of IPA (QIAGEN Inc., https://www.qiagenbioinformatics.com/products/ingenuity-pathway-analysis). To identify previously annotated ligands, we crossed referenced our gene list with the FANTOM5 database (fantom.gsc.riken.jp/5/)^3^. To assess the enrichment for ligands in p16+ mesenchyme, we calculated the statistical significance of the overlap (ligands identified to be upregulated in p16+ mesenchyme) between total identified ligands in FANTOM5 (gene set 1) and total number of upregulated genes (gene set 2) utilizing the webtool on nemate.org, which calculates the representation factor along with the p value of overlap as a measure of the significance of enrichment.

### Cell Culture

Freshly isolated PDGFRα+ cells from *INBRITE* lungs (GFP- or GFP+) were cultured on gelatin-treated tissue culture plates in DMEM-F12 with 10% FBS and 1% Penicillin-Streptomycin. Media was refreshed every other day and primary lung mesenchymal cells were maintained for no more than five passages. For the WST1 proliferation assay, cells were plated in a 96 well plate at a concentration of 1000 cells/well for 72 hours to allow adhering. After cells adhered the first data point was read as time zero by adding 1:10 of cell proliferation Reagent WST-1 (Sigma, #11644807001) to cells and incubating them at 37 °C for 30 minutes. Each plate was read using Biotek H1 plate reader at 420-480nm with max absorption at 440nm. For cell size assessment, the cells were fixed on the plate and stained with DAPI and F-actin with phalloidin-555 (Thermo, #34055) for 1 hour at room temperature. Phalloidin was used at (125 U/ml) concentration. Cell size was measured by using the freehand line feature to outline each cell from 40X magnification images and Measure to get area. All cells in GFP- or GFP+ mesenchyme wells were assayed and averaged and compared for statistical analysis.

### Cytospin

Freshly sorted cells were rinsed in 1X PBS for 5 minutes spun down at 550g for 5 minutes then fixed with 4% PFA for 15 minutes at room temperature. After fixation cells were rinse with 1X PBS, centrifuged down to get rid of supernatant, and then resuspended in 50μl of fresh 1X PBS. Cells were spun onto superfrost slides at 340g for 5 minutes. Stain for F-actin and DAPI was performed as described above.

### Organoid Assay

For organoid assay, *INKBRITE* adult lungs were FACS sorted for CD45-Epcam-CD31-PDGFRa+GFP- or GFP+ mesenchymal cells and epithelial cells were sorted from *Scgb1a1*^*creERT2*^*:R*^*tdt*^ tamoxifen induced animals. Scgb1a1+ epithelial cells and GFP- or GFP+ mesenchymal cells were co-cultured (1.5×10^4^ epithelial cells : 5×10^4^ mesenchymal cells/well) in a modified MTEC media diluted 1:1 in growth factor reduced matrigel. Modified MTEC culture media is comprised of small airway basal media (SABM) with selected components from SAGM bullet kit (Lonza) including Insulin, Transferrin, Bovine Pituitary Extract, Retinoic Acid, and human Epidermal Growth Factor. 0.1 μg/mL cholera toxin, 5% FBS, and 1% Penicillin-Streptomycin were also added. Cell suspension-matrigel mixture was placed in a transwell and incubated in growth media with 10 μM ROCK inhibitor in a 24 well plate for 48 hours, after which the media was replenished every other day (lacking ROCK inhibitor). Each experimental condition was performed in triplicates. Where applicable, recombinant Ereg (2 ng/mL, R&D, #1068) and afatinib (5 μM, Selleckem, #1011) were added to the media after 48 hours and replenished in every media change. Colonies were assayed after 12-14 days. Each transwell was imaged using EVOS M5000 at 1.25X magnification and quantified on ImageJ blind to experimental condition.

### Quantitative RT-PCR

Total RNA was isolated from epithelial cells isolated from organoid assays or cultured primary lung fibroblasts using the PicoPure RNA Isolation Kit or the RNeasy Kit, following the manufacturers’ protocols. RNA from mouse lung tissue was obtained by removing the entire left lobe or from sorted single cell suspension, homogenizing in trizol, and extracting using the E.Z.N.A Total RNA Kit following manufacturer instructions. cDNA was synthesized from total RNA using the SuperScript Strand Synthesis System. Quantitative PCR was performed using the SYBR Green system. Primers are listed in below. Relative gene expression levels after qRT-PCR were defined using the ΔΔCt method and normalizing to GAPDH. We used a minimum of three biological replicates for each genotype/condition. One-tailed t tests were used to perform statistical analysis of fold changes between genotypes/conditions.

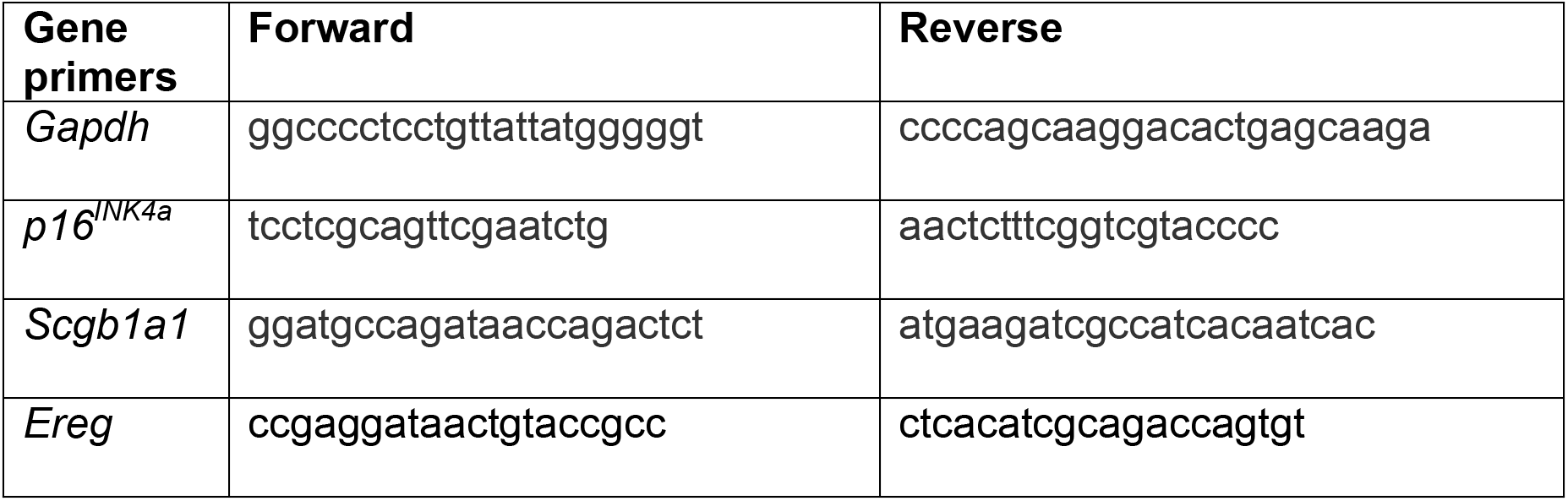

### Immunofluorescence Image Quantification

Sections were imaged for quantification on a Zeiss Lumar V12 microscope. At least three samples per genotype/condition were used, and at least 5-10 randomly selected sections were chosen for each sample. Cell counts for SGCGB1A1+, Brdu+, GFP+, RFP+, and DAPI cells were performed on Fiji using the “Cell Counter” plug-in. For quantification of airway length the freehand line feature couple with Measure. Cell size was measured by using the freehand line feature to outline each cell from 40X magnification images and Measure to get area. All cells per GFP- or GFP+ mesenchyme were averaged and compared for statistical analysis.

### Statistical Analysis

All statistical analyses were performed in GraphPad Prism 6.0. One-tailed t-tests were used to determine the p value and data in graphs are presented as mean ± SEM.

### Data and Software Availability

The sequencing data reported in this paper is deposited in NCBI Gene Expression Omnibus (GEO) under the accession number GSE140654. The Gli1+ single cell sequencing data is deposited separately under the accession number GSE140032.

## Acknowledgements

We thank Dean Sheppard for critical review of manuscript; Ned Sharpless for providing the p16 flox allele; the Parnassus Flow Cytometry Core for assistance with cell sorting for bulk and single cell RNA analysis (P30DK063720); Biological Imaging Development Core members (P30 DK063720); Eunice Wan and the Institute for Human Genetics Core for processing of single cell RNA samples and high-throughput sequencing. GEO accession number for raw RNA sequencing data is listed in Materials and Methods. This work is supported by NIH grants DP2AG056034, K08HL121146, R01HL142552 to T.P., along with Tobacco Related Disease Research Program New Investigator Award and Pulmonary Hypertension Association award to T.P. and F32HL14226 to N.R.

## Author Contributions

N.R. and T.P. conceived the experiments. N.R., K.C., M.C., C.W., P.M., and M.K. performed the experiments and data analysis. A.M., and J.C. provided expertise and feedback. N.R. and T.P. wrote the manuscript.

## Competing Interest Declaration

None

**Supplementary Information** is available for this paper.

Correspondence and requests for materials should be addressed to Tien Peng.

## Notes

### Competing Interest Statement

The authors have declared no competing interest.

## Reference

1 van Deursen, J. M. Senolytic therapies for healthy longevity. Science 364, 636–637, doi:10.1126/science.aaw1299 (2019).

2 Sharpless, N. E. & Sherr, C. J. Forging a signature of in vivo senescence. Nat Rev Cancer 15, 397–408, doi:10.1038/nrc3960 (2015).

3 Lopez-Otin, C., Blasco, M. A., Partridge, L., Serrano, M. & Kroemer, G. The hallmarks of aging. Cell 153, 1194–1217, doi:10.1016/j.cell.2013.05.039 (2013).

4 Gorgoulis, V. et al. Cellular Senescence: Defining a Path Forward. Cell 179, 813–827, doi:10.1016/j.cell.2019.10.005 (2019).

5 Baker, D. J. et al. Clearance of p16Ink4a-positive senescent cells delays ageing-associated disorders. Nature 479, 232–236, doi:10.1038/nature10600 (2011).

6 Baker, D. J. et al. Naturally occurring p16(Ink4a)-positive cells shorten healthy lifespan. Nature 530, 184–189, doi:10.1038/nature16932 (2016).

7 Chang, J. et al. Clearance of senescent cells by ABT263 rejuvenates aged hematopoietic stem cells in mice. Nat Med 22, 78–83, doi:10.1038/nm.4010 (2016).

8 Childs, B. G. et al. Senescent intimal foam cells are deleterious at all stages of atherosclerosis. Science 354, 472–477, doi:10.1126/science.aaf6659 (2016).

9 Baar, M. P. et al. Targeted Apoptosis of Senescent Cells Restores Tissue Homeostasis in Response to Chemotoxicity and Aging. Cell 169, 132–147 e116, doi:10.1016/j.cell.2017.02.031 (2017).

10 Jeon, O. H. et al. Local clearance of senescent cells attenuates the development of post-traumatic osteoarthritis and creates a pro-regenerative environment. Nat Med 23, 775–781, doi:10.1038/nm.4324 (2017).

11 Bussian, T. J. et al. Clearance of senescent glial cells prevents tau-dependent pathology and cognitive decline. Nature 562, 578–582, doi:10.1038/s41586-018-0543-y (2018).

12 Liu, J. Y. et al. Cells exhibiting strong p16 (INK4a) promoter activation in vivo display features of senescence. Proc Natl Acad Sci U S A 116, 2603–2611, doi:10.1073/pnas.1818313116 (2019).

13 Zindy, F., Quelle, D. E., Roussel, M. F. & Sherr, C. J. Expression of the p16INK4a tumor suppressor versus other INK4 family members during mouse development and aging. Oncogene 15, 203–211, doi:10.1038/sj.onc.1201178 (1997).

14 Takeuchi, S. et al. Intrinsic cooperation between p16INK4a and p21Waf1/Cip1 in the onset of cellular senescence and tumor suppression in vivo. Cancer Res 70, 9381–9390, doi:10.1158/0008-5472.CAN-10-0801 (2010).

15 Burd, C. E. et al. Monitoring tumorigenesis and senescence in vivo with a p16(INK4a)-luciferase model. Cell 152, 340–351, doi:10.1016/j.cell.2012.12.010 (2013).

16 Demaria, M. et al. An essential role for senescent cells in optimal wound healing through secretion of PDGF-AA. Dev Cell 31, 722–733, doi:10.1016/j.devcel.2014.11.012 (2014).

17 Hong, K. U., Reynolds, S. D., Giangreco, A., Hurley, C. M. & Stripp, B. R. Clara cell secretory protein-expressing cells of the airway neuroepithelial body microenvironment include a label-retaining subset and are critical for epithelial renewal after progenitor cell depletion. Am J Respir Cell Mol Biol 24, 671–681, doi:10.1165/ajrcmb.24.6.4498 (2001).

18 Peng, T. et al. Hedgehog actively maintains adult lung quiescence and regulates repair and regeneration. Nature 526, 578–582, doi:10.1038/nature14984 (2015).

19 Munoz-Espin, D. et al. Programmed cell senescence during mammalian embryonic development. Cell 155, 1104–1118, doi:10.1016/j.cell.2013.10.019 (2013).

20 Storer, M. et al. Senescence is a developmental mechanism that contributes to embryonic growth and patterning. Cell 155, 1119–1130, doi:10.1016/j.cell.2013.10.041 (2013).

21 Benn, P. A. Specific chromosome aberrations in senescent fibroblast cell lines derived from human embryos. Am J Hum Genet 28, 465–473 (1976).

22 Yamamoto, M., Mitsui, Y., Ooka, H. & Yamamoto, K. Appearance of the terminal senescent cell population in human diploid fibroblasts analyzed by flow cytometry. Mech Ageing Dev 51, 195–214, doi:10.1016/0047-6374(90)90071-m (1990).

23 Zhu, Y. Z. et al. Hepatitis B virus X protein promotes hypermethylation of p16(INK4A) promoter through upregulation of DNA methyltransferases in hepatocarcinogenesis. Exp Mol Pathol 89, 268–275, doi:10.1016/j.yexmp.2010.06.013 (2010).

24 Dimri, M., Carroll, J. D., Cho, J. H. & Dimri, G. P. microRNA-141 regulates BMI1 expression and induces senescence in human diploid fibroblasts. Cell Cycle 12, 3537–3546, doi:10.4161/cc.26592 (2013).

25 Chen, Z. et al. miR-124 and miR-506 inhibit colorectal cancer progression by targeting DNMT3B and DNMT1. Oncotarget 6, 38139–38150, doi:10.18632/oncotarget.5709 (2015).

26 Chang, S. et al. Hypoxic reprograming of H3K27me3 and H3K4me3 at the INK4A locus. FEBS Lett 590, 3407–3415, doi:10.1002/1873-3468.12375 (2016).

27 Pandey, R. et al. MicroRNA-1825 induces proliferation of adult cardiomyocytes and promotes cardiac regeneration post ischemic injury. Am J Transl Res 9, 3120–3137 (2017).

28 Laberge, R. M. et al. MTOR regulates the pro-tumorigenic senescence-associated secretory phenotype by promoting IL1A translation. Nat Cell Biol 17, 1049–1061, doi:10.1038/ncb3195 (2015).

29 Ramilowski, J. A. et al. A draft network of ligand-receptor-mediated multicellular signalling in human. Nat Commun 6, 7866, doi:10.1038/ncomms8866 (2015).

30 Toyoda, H. et al. Epiregulin. A novel epidermal growth factor with mitogenic activity for rat primary hepatocytes. J Biol Chem 270, 7495–7500, doi:10.1074/jbc.270.13.7495 (1995).

31 Draper, B. K., Komurasaki, T., Davidson, M. K. & Nanney, L. B. Epiregulin is more potent than EGF or TGFalpha in promoting in vitro wound closure due to enhanced ERK/MAPK activation. J Cell Biochem 89, 1126–1137, doi:10.1002/jcb.10584 (2003).

32 Li, S. et al. Mesenchymal-epithelial interactions involving epiregulin in tuberous sclerosis complex hamartomas. Proc Natl Acad Sci U S A 105, 3539–3544, doi:10.1073/pnas.0712397105 (2008).

33 Sosic, D., Richardson, J. A., Yu, K., Ornitz, D. M. & Olson, E. N. Twist regulates cytokine gene expression through a negative feedback loop that represses NF-kappaB activity. Cell 112, 169–180, doi:10.1016/s0092-8674(03)00002-3 (2003).

34 Kramann, R. et al. Perivascular Gli1+ progenitors are key contributors to injury-induced organ fibrosis. Cell Stem Cell 16, 51–66, doi:10.1016/j.stem.2014.11.004 (2015).

35 Dahlgren, M. W. et al. Adventitial Stromal Cells Define Group 2 Innate Lymphoid Cell Tissue Niches. Immunity 50, 707–722 e706, doi:10.1016/j.immuni.2019.02.002 (2019).

36 Wang, C. et al. Expansion of hedgehog disrupts mesenchymal identity and induces emphysema phenotype. J Clin Invest 128, 4343–4358, doi:10.1172/JCI99435 (2018).

37 Scudellari, M. To stay young, kill zombie cells. Nature 550, 448–450, doi:10.1038/550448a (2017).

38 Dimri, G. P. et al. A biomarker that identifies senescent human cells in culture and in aging skin in vivo. Proc Natl Acad Sci U S A 92, 9363–9367, doi:10.1073/pnas.92.20.9363 (1995).

## Method Reference

1 Demaria, M. et al. An essential role for senescent cells in optimal wound healing through secretion of PDGF-AA. Dev Cell 31, 722–733, doi:10.1016/j.devcel.2014.11.012 (2014).

2 Wang, C. et al. Expansion of hedgehog disrupts mesenchymal identity and induces emphysema phenotype. J Clin Invest 128, 4343–4358, doi:10.1172/JCI99435 (2018).

3 Ramilowski, J. A. et al. A draft network of ligand-receptor-mediated multicellular signalling in human. Nat Commun 6, 7866, doi:10.1038/ncomms8866 (2015).

